# Spatial scaling of beta diversity supports the regional community concept for clades as different as ants, birds, diatoms, and trees

**DOI:** 10.1101/2023.10.24.563827

**Authors:** Leo Ohyama, Juan D. Bogota-Gregory, David G. Jenkins

## Abstract

**Aim:** Three fundamental and inter-related concepts have accrued debates: ecological communities, beta diversity (β), and spatial scale. Spatial scaling of β informs the community concept because the scale of maximal β corresponds to the most apparent size of an ecological community (without invoking external features of habitat, etc.). Here we test five alternative hypotheses about spatial scaling of β for ants, birds, diatoms, and trees across the contiguous USA, using spatial grains from 1 to 10^6^ km^2^. We compare β scaling among clades and test hypotheses about repeatability where data permit for: (a) summer and winter bird β in six consecutive years; (b) trees through time (4 years, spaced 5 years apart). Finally, we compare different forms of β (i.e., observed and deviations from null models based on spatial heterogeneity and spatial homogeneity).

**Location:** The contiguous United States of America

**Time Period:** Recent but varying with clade

**Taxa Studied:** ants, birds, diatoms, and trees

**Methods:** We obtained data from publicly-available sources and assigned point locations to hexagonal grids ranging from 1 to 10^6^ km^2^. At each spatial grain, we calculated mean pairwise β between each hexagon and its neighboring grids. We also compared alternative β measures and evaluated potential confounding effects of neighborhood size and species richness on results.

**Results:** Spatial scaling of β repeatedly supported the regional community concept among clades, though with different spatial scales per clade. Based on peak mean β, community size for trees (∼300 km^2^) < winter birds (∼500 km^2^) < summer birds (∼2000 km^2^) ≈ ants (∼2000 km^2^) < diatoms (∼11,000 km^2^). We note that community scales represent peaks on gradients rather than definitive one-size-fits-all scales. Spatial scaling of β was sensitive to seasonality (birds) and consistent among years for both birds and trees. Also, β deviation from a null model based on spatial heterogeneity adjusted observed β but was less sensitive to neighborhood size and species richness than β deviation based on spatial homogeneity.

**Main conclusions:** Results here indicate that: (a) similar patterns should occur across the tree of life; (b) local ecological and evolutionary forces scale up to form repeatable regional community patterns in ways not yet fully understood; (c) local biodiversity conservation efforts need to be coordinated at biogeographical scales to best achieve goals; and (d) a recent method to calculate β deviation from a null model based on spatial heterogeneity improves β research.

## Introduction

Ecological communities are often defined as “multiple species living in a specified place and time” (Vellend 2010) and are a common and important focal point in ecology. This definition is purposefully flexible in its spatial scale (i.e., grain and extent; Wiens 1989) to include different study systems and taxa. However, it leaves hanging a question at the heart of critiques for ecology (McIntosh 1986, Wilson 1991, Lawton 1999, Ricklefs 2008, Roughgarden 2009, Vellend 2010): Are ecological communities real, natural objects or merely abstract ideas?

While this question is fundamental with important practical applications, the lack of a clear answer has obstructed progress for a century. Ecologists have navigated around this uncertainty when studying natural systems often by one of four methods. A common method is to use an operational definition (e.g., the one above) to bear “as little conceptual baggage as possible, so that we can put the debate about their existence behind us” (Palmer and White 1994). Some disagreed with that method (Wilson 1994, Jax et al. 1998), leading to reliance instead on external proxies (i.e., not based on community patterns) such as geomorphology or habitat (Cadenasso et al. 2003, Fagan et al. 2003, Yarrow and Marin 2007). A third method is to focus on levels of organization that lie below (i.e., individual organisms or populations; Clark et al. 2011) or above (i.e., metacommunities; Leibold et al. 2004) the community level.

Unfortunately the fundamental problem remains as such methods like metacommunity ecology still rely on communities as their fundamental unit. Finally, Ricklefs (2008) argued that the problem is one of scale, and that a focus on local communities should be abandoned in favor of regional scales to better address the full suite of selective conditions driving diversity.

Beyond basic ecology, the veracity of ecological communities is relevant to studying Anthropocenic biodiversity and conservation biology. Major changes to biodiversity may occur due to the interaction between land use and climate change, but detection and prediction of those effects likely depend on the scale at which changes are studied. Also, the efficacy of conservation efforts, including coordinated corridors and networks among conserved sites (e.g., Rouget et al. 2006) may depend in part on the spatial scale of those efforts. Even after all this time and debate, it appears that research on the scaling of biodiversity patterns may help resolve this lingering uncertainty on the ecological communities, with potential applications to better understand the magnitude and direction of the effects of ongoing anthropogenic activities and changes (e.g., land use and climate change) on biodiversity.

Here we answer the deceptively simple question “how big is a community?” by evaluating beta diversity, which represents the difference between two communities (Whittaker 1972, Tuomisto 2010, Anderson et al. 2011). Hereafter we denote beta diversity in the general sense with β, where specific versions are denoted by subscripts below. We adapt long-standing logic from research on spatial resolution in remote sensing (e.g., Woodcock and Strahler 1987, Ming et al. 2011, Mutowo et al. 2023) to reason that maximal β indicates the maximal resolution between neighboring communities, and so the matching spatial resolution corresponds to the most-apparent, empirical size of a community. We note that this approach relies on the focal organisms to indicate patterns; it does not make assumptions based on external landscape or habitat features, nor does it make assumptions based on population or metapopulation biology (e.g., dispersal distances, etc.). Additionally, our approach obtains a quantitative *gradient* showing maximal β and associated variance at some spatial scale; as such, it does not presume thresholds or categories. To more fully understand effects of external conditions or intrinsic traits is to undertake a different, and important body of research; our research simply seeks to identify spatial scales at which that research may best reflect community patterns. We work with data for four very different clades (ants, birds, diatoms, and trees) to consider potential generality of pattern; thus, the community concept is represented here as assemblages (Fauth et al. 1996, Stroud et al. 2015).

We tread carefully with both β and spatial scale, which are entangled throughout the literature and require further explanation below. Beta diversity bears enough confusion accrued from the past decades to be described as “a concept gone awry” (Tuomisto 2010) with a “confusing array of concepts, measures and methods” (Anderson et al. 2011). Here we use a β based on a multiplicative relationship: β = γ/α, where γ is overall diversity and α is local diversity (Tuomisto 2010; Xing & He 2021)). An even more important decision for our purposes here is to use a deviation of β from randomized expectations (e.g., β_DEV_). This matters because β is a function of both γ and α. Thus β in neighborhoods of sites with different γ and/or α will inherently differ but can be adjusted to more fairly evaluate spatial patterns by calculating the difference from randomizations conducted with the same data. Initially, β_DEV_ was calculated using a spatially-homogeneous null (Kraft et al. 2011), but a recent version based on spatial heterogeneity (called β_NBD_ for its negative binomial distribution) can represent patterns more accurately across spatially-varying regions (Xing & He 2021). Here we use β_NBD_ as our main measure of β but also compare results to β_DEV_ and observed β (β_OBS_).

Spatial scale is important to ecology because both patterns (e.g., β) and processes have inherent scales (Wiens 1989, Barton et al. 2013, Zhang et al. 2015); to discern well the process-pattern linkage requires matching scales. Here we hold spatial extent constant (the contiguous USA; 8.08 x 10^6^ km^2^) to evaluate the relationship between β and spatial grain (i.e., resolution) across a million-fold range (1 km^2^ to 1 million km^2^).

Like other measures in ecology, β is expected to depend on spatial grain in at least five different ways (Fig. 1), where that variety appears related (in part) to the variety of β measures and spatial scales that have been studied. If maximal β occurs at small spatial grains (Fig. 1a), then locally-scaled communities would be supported (Barton et al. 2013, Keil et al. 2012).

**Fig. 1.**
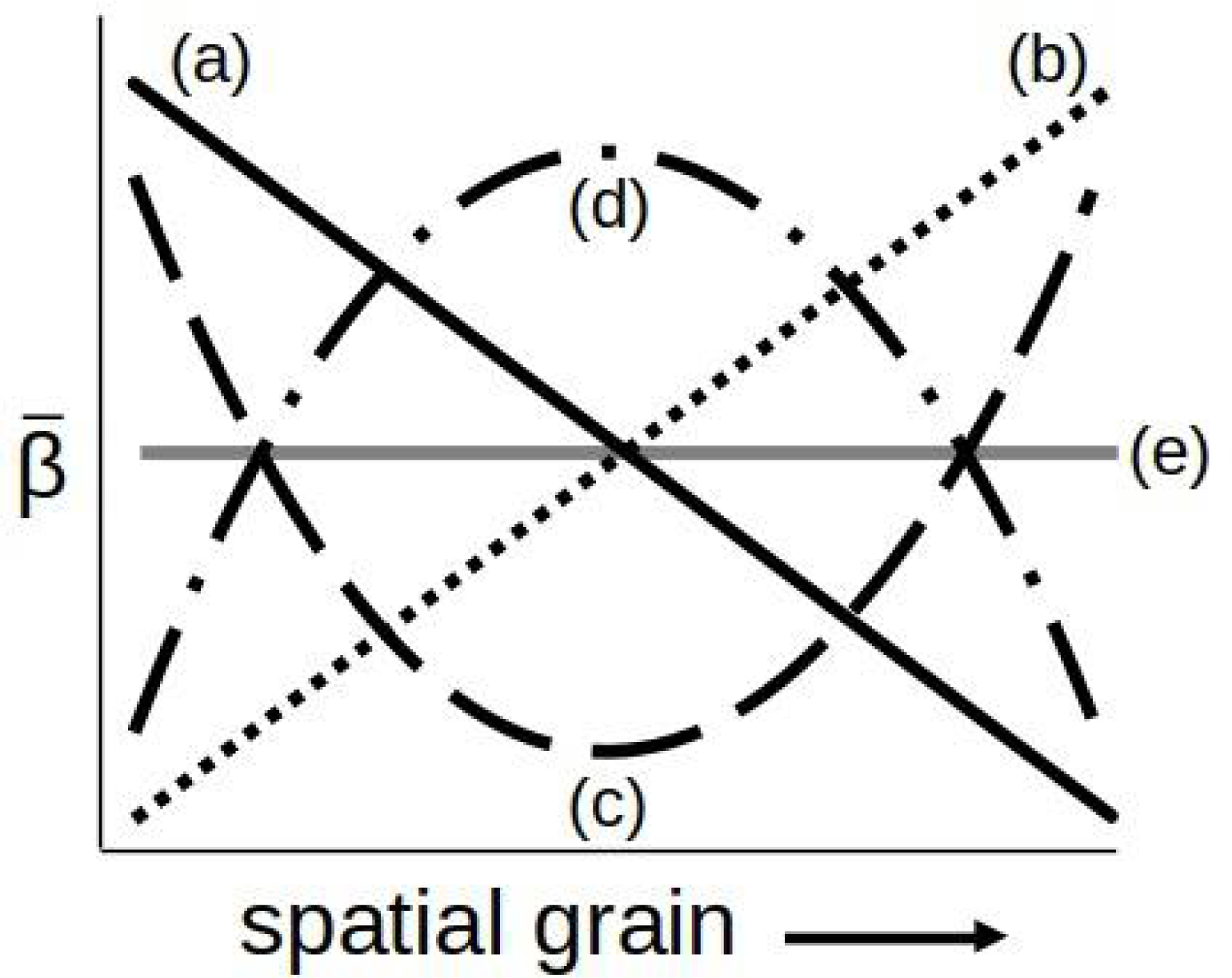
Hypothetical shapes of beta diversity spatial scaling. As spatial grain increases, beta diversity may (a) decrease because diversity patterns are caused by local processes (e.g., Barton et al. 2013, Keil et al. 2012); (b) increase because more samples at greater grain increase beta diversity (e.g., Zhang et al. 2015, Keil and Chase 2019, Xing and He 2021); (c) first decrease and then increase because both effects in (a) and (b) occur or have been studied at different scales; (d) peak at intermediate scales because diversity patterns are caused by regional processes (Ricklefs 2008); or (e) be uninformative (e.g., Lira-Noriega et al. 2007).

Alternatively, β may progressively increase with spatial grain, reflecting sample size effects (Fig 1b; Zhang et al. 2015, Keil and Chase 2019, Xing and He 2021). Both trends may occur but represent studies conducted at different spatial scales, so β may first decrease and then increase to follow a V- or U-shaped curve across a wider range of spatial grains (Fig. 1c). Alternatively, the regional community concept (Ricklefs 2008) would be supported if β exhibited hump-shaped peaks at an intermediate spatial grain (Fig. 1d), which would also be consistent with patterns of an optimized spatial grain in remote sensing (Woodcock and Strahler 1987, Ming et al. 2011, Mutowo et al. 2023). Finally, it is possible that β exhibits no clear trend with spatial grain (Fig. 1e; Lira-Noriega et al. 2007).

If spatial scale is composed of two axes (grain and extent), a study scale represents a third axis, often measured as sample size (e.g., Jenkins 2011, Xing & He 2020). Study scale can affect observed β if larger areas include more samples that enable the detection of more species, as expressed in rarefaction of species-sample curves (Chao et al. 2014). We addressed study scale effects across spatial grains using both *a priori* and *a posteriori* approaches. We pooled separate, georeferenced samples into each geographic grid cell (where size of the cell represents spatial grain) to estimate diversity in a cell’s area. We then compare each grid cell to each of its neighbors, where hexagonal grids are constrained to have ≤ 6 neighbors (thus controlling grid sample size across spatial grains). Pairwise comparisons of grid cells (i.e., β) avoids errors inherent in other methods (Marion et al. 2017). A tradeoff may still linger, where smaller grid cells contain too few samples for reliable diversity estimation (i.e., errors of omission) but very large grid cells are so inclusive as to blur actual patterns (i.e., errors of commission; Boschetti et al. 2004). Thus we also evaluate *a posteriori* any lingering effects of sample size per se (i.e., the number of data points within a hexagonal grid) on results across spatial grains. We also evaluate effects of neighborhood size (M) and species richness (S) on β measures to show that the use of β_NBD_ is supported, whereas β_DEV_ is problematic (Supplementary Information).

In summary, we control scaling effects of spatial extent and sample size to evaluate potential β ∼ spatial grain relationships. We then evaluate mean pairwise β (in several forms) among spatial hexagons ranging from 1.0 to 1 million km^2^ for four very different clades (ants, birds, diatoms, and trees). Because data sets differed substantially in their sources, subject taxa, and features (Table 1), we reasoned that similar β scaling shape (i.e., a line in Fig. 1) across datasets could indicate generality for varied taxa and data origins. However, we expected data sets to differ in details of β scaling.

**Table 1.** Summaries of analyzed data sets.

|  | Ants | Birds | Diatoms | Trees |
| --- | --- | --- | --- | --- |
| Number of species | 699 | 1,560 | 1,957 | 403 |
| Number of samples | 24,893 | 810,921 | 1,070 | 39,396 |
| Temporal range | 2 centuries | 2010 – 2015 | 2007 | 2002-2017* |
| Data type | presence | Mean count per<br>person-hour | presence | presence |
| Temporal analyses? | No | Yes; seasonal | No | Yes; annual |
| Data Source | Guénard et al.<br>(2017) | Sullivan et al.<br>(2009) | USEPA<br>(2009) | USDA<br>(2019) |

\* 5-year increments

Data for birds and trees enabled comparisons within each data set; we analyzed bird data for winter and summer in each of six consecutive years to evaluate sensitivity of this approach to seasonal changes and interannual repeatability of those patterns. We hypothesized that seasonal migrations would concentrate winter birds in the southern USA compared to summer to cause reduced: (a) spatial scale of bird communities (i.e, the grain at which peak β_NBD_ occurs); and (b) magnitude of peak β_NBD_. We also expected annual re-assembly cycles in bird communities would result in repeatable spatial scaling and patterns during the 6-year span. Similarly, we hypothesized similar β spatial scaling of trees during 15 years despite varied sample locations each year.

Beyond β_NBD_ scaling shape (Fig. 1), differences in the details among clades could inform long-standing questions of dispersal scaling of diversity among microbial and macroscopic organisms (Fontaneto 2011). Assuming similar general β_NBD_ spatial scaling shapes across clades, we expected diatoms to exhibit smaller spatial grains for peak β_NBD_ than other taxa because: diatoms exist within hard ecotones (i.e., lake shores here); are sensitive to local water quality (Reid et al. 1995); and most species inhabit one or few lakes (USEPA 2009). In comparison, species of trees, ants and birds inhabit intergrading habitats, have aerial life stages, and are known to inhabit substantial ranges. In combination with the general scaling of dispersal distances for actively dispersing organisms (Jenkins et al. 2007), we naively expected spatial scales of peak β_NBD_ would rank as diatoms < ants < trees < birds.

## Methods

We obtained data for the contiguous USA for ants (Guénard et al. 2017), birds (Sullivan et al. 2009), diatoms (USEPA 2009), and trees (USDA Forest Service 2019; Table 1). Ant data represented cumulative records synthesized from literature records and existing databases. Any ant observations marked as dubious or needing confirmation were removed prior to analysis. The eBird system (Sullivan et al. 2009) amasses standardized community science data, including the number of birds per species, number of observers and time spent. Bird data analyzed here represented cleaned reports from January and July during 2010 - 2015. We expressed bird abundances per unit effort (i.e., per person-hours) to standardize among reports. We used January to represent winter conditions and July to represent summer conditions. Diatom data analyzed here were collected during one year from the top 5 cm of lake sediment and processed according to standardized protocols (USEPA 2009).

Data for trees (i.e., stems with diameter at breast height > 5 cm) for the years 2000-2020 were extracted from the US Forest Service’s Forest Inventory Analysis (FIA) data using the rFIA package in R (Stanke et al. 2020, R Core Team 2021). The years 2002, 2007, 2012, and 2017 were retained for analyses here because they had similar numbers of samples and were evenly separated in time. More information on FIA’s standardized sampling protocols is available at https://www.fia.fs.usda.gov/library. Geographical locations of forest sample data in FIA are “fuzzed” by <1 mile and 20% are “swapped” with another similar, nearby property to shield private lands in the publicly-available data (Burrill et al. 2021). Accordingly, we truncated the lower limit for overall mean β spatial scaling of trees to be 10-fold greater than other data sets (i.e., 10 km^2^).

Each taxon was analyzed similarly using an iterative process summarized here, based on multiple R packages (R Core Team 2021; see code in Supplementary Information). Georeferenced data were cleaned for uncertain species identities and intersected with hexagonal grids. Where data represented counts (birds and trees), one datum per species per hexagon was obtained by averaging point data. Bird counts were standardized as mean count per person-hour, rounded up to the nearest whole number. Data for ants and diatoms were presence/absence; a species present somewhere in a hexagonal grid was listed as present. To be clear, β here was based on species presence/absence in an area, but β_NBD_ calculated null models and thus deviations using abundances, if available (Xing and He 2021).

Hexagonal grids and their 1^st^-degree adjoining neighbors comprised a neighborhood. All β calculations were conducted between a central grid and each of its neighbors, where a lower limit for calculations was that a central grid had at least two sites and 3 species. Finest spatial grains were most likely affected by this threshold. This process was repeated at spatial grains (i.e. hexagonal grid areas) from 1 km^2^ to 1 million km^2^ (i.e., 1, 5, 10, 50, … etc.; 13 spatial grains in total). Mean pairwise β for each hexagon enabled fair comparisons among hexagons within and among spatial grains.

Because data sets differed (Table 1), β was calculated for birds and trees at more times (two seasons in six consecutive years for birds, four years for trees) than for ants or diatoms. In addition, we calculated multiple β versions per Xing & He (2020); we emphasize β_NBD_ for reasons explained above, but also present in Supplementary Information the results for observed β (β_OBS_; Tuomisto 2010) and the spatially-homogeneous deviation (β_DEV_; Kraft et al. 2011). Observed β is Tuomisto’s (2010) proportional species turnover (i.e., ^0^β_Pt_ = 1 − α ⁄ γ, where α = mean number of species in a hexagon and γ = total number of species in the pair of hexagons being compared; *18*).

We evaluated alternative hypotheses (Fig. 1) graphically in two ways. We used general additive models to fit curves to data, where each point represented a mean β_NBD_ (± 95% confidence intervals) at a spatial grain size. We also evaluated β_NBD_ distributions at each grain size to represent variation around means. Graphical results were generated using ggplot2 and ggridges in R (Wickham 2016, Wilke 2022). We also ensured correct interpretation relative to hypotheses (Fig. 1) by comparing linear and quadratic models of data, using Akaike Information Criteria weights (computed with bbmle in R; Bolker et al. 2022), corrected for sample size (i.e., AICc *w_i_*), which represent the probability that a model is most efficient among those compared; Burnham and Anderson 2002). Model coefficient signs for the most plausible model then identified curve direction. Maps of mean pairwise β_NBD_ informed interpretations, and comparisons of β_NBD_ to β_DEV_ and β_OBS_ confirmed the results of Xing & He (2020), including the value of β_NBD_ for analyses.

We also evaluated potential sample size effects on β_NBD_ results, because β_DEV_ increases with greater sample size but such an effect was undescribed for β_NBD_ by Xing & He (2021). Because sample size can be measured in different ways, we examined sample size as: simple sample size (i.e., number of sample points in hexagons), number of hexagons (that contribute mean β_NBD_), neighborhood size (constrained for hexagons to ≤6), and species richness.

## Results

Ants, birds, diatoms, and trees all demonstrated peak β_NBD_ at intermediate spatial grains that were roughly consistent in shape (Fig. 2), despite wide differences in natural history and data (Table 1). As may be expected for ants, birds, and diatoms from Fig. 2, hump-shaped curves more efficiently represented data than linear models that were consistent with other scaling hypotheses (all AICc weights > 0.95; Fig. 1). Moreover, curve coefficients supported hump-shaped curves rather than V- or U shaped curves (again, see Fig. 1). However, the variable but increased peak β_NBD_ at greatest spatial grains for trees rendered null models most plausible. The impact of grains = 500,000 and 1 million km2 on that analysis was demonstrated by omitting them from analyses; again hump-shaped patterns were confirmed (AICc weights > 0.95). The broadly consistent result across very different clades and data sets was consistent with the regional community concept. Moreover, interannual or interseason variation (where possible; birds, trees) indicated remarkable consistency.

**Fig. 2.**
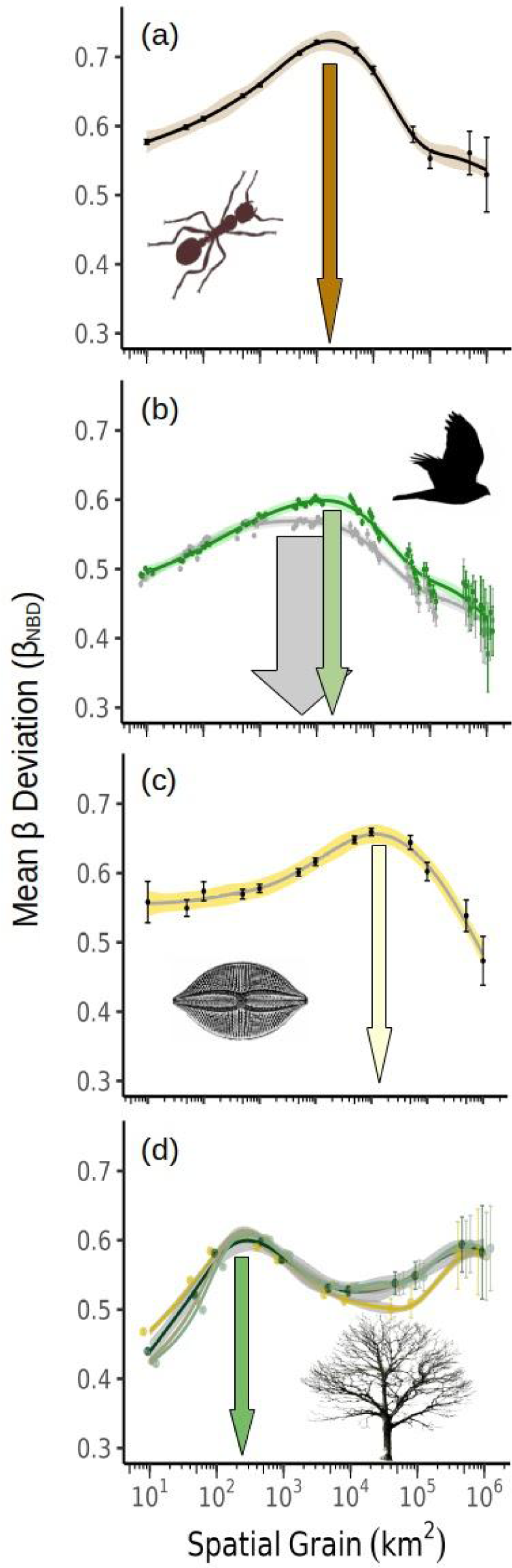
Spatial scaling of beta diversity, calculated as deviation from a null expectation (β_DEV_, *29*) and potentially ranging 0-1. β_NBD_ scaling of (a) ants; (b) birds (summer = green; winter =grey, 2010-2015); (c) diatoms; and (d) trees (2002, 2007, 2012, and 2017). Arrows indicate spatial grains corresponding to approximate peak β_DEV_. Note log scale for spatial grain. Public domain organism images.

Within that broad pattern, values of peak β_NBD_ (Fig. 2) differed substantially among clades, and our hypothesis about the rankings of β_NBD_ was incorrect. Peak mean β_NBD_ for ants was about 1,500 km^2^ (Fig. 2a). Peak mean β_NBD_ values for birds were about ∼ 2,000 km^2^ in summer but substantially reduced to roughly 500 km^2^ in winter (green and gray vertical arrows in Fig. 2b, respectively). Peak mean β_NBD_ values were greatest for diatoms (∼11,000 km^2^; Fig. 2c), but for trees peak mean β_NBD_ values were consistently at about 300 km^2^ through 4 evaluated years spanning 2 decades (Fig. 2d). Unlike other clades, mean β_NBD_ for trees again increased at greatest spatial grains (Fig. 2d). However, values at about 300 km^2^ had very little within- or among-annual variation (95% confidence intervals are smaller than symbols), whereas the values at greatest spatial grain had wide confidence intervals (Fig. 2b), for reasons we explain below and which led us to disregard the second peak. Thus, we ranked peak mean β_NBD_ among clades as [trees < winter birds < ants ≈ summer birds < diatoms], rather different from our naive hypothesis (diatoms < ants < trees < birds).

We also evaluated variation of β_NBD_ as a function of spatial grain, where each central hexagon’s mean β_NBD_ contributed to a plotted distribution (note the “flipped” axes between Figs. 2 & 3). Essentially, wider confidence intervals around peak mean β_NBD_ (Fig. 2) were related to the number of hexagons contributing to a mean (e.g., only 14 at 1 million km^2^ grain) and variation in mean β_NBD_ values (Fig. 3). Ants had more variation in β_NBD_ at any one spatial grain than other clades (Fig. 3a), and multimodal distributions of β_NBD_ were apparent at both smallest and greatest grains, while unimodal and narrower distributions at peak β_NBD_ were most appropriate for a mean (Fig. 3a). Distributions of β_NBD_ were narrow for birds, so that mean values in Fig. 2 represented patterns quite well regardless of season, and especially at spatial grains < 500,000 km^2^ (Figs. 3b & 3c). Like ants, distributions for diatoms were multimodal at smallest and greatest grains but appropriate to a mean value at intermediate grains where peak values occurred (Fig. 3d). Distributions for trees supported the use of mean peak β_NBD_ values at smaller grains, but variation at greater grains reduced validity of a mean to represent patterns (Fig. 3e).

**Fig. 3.**
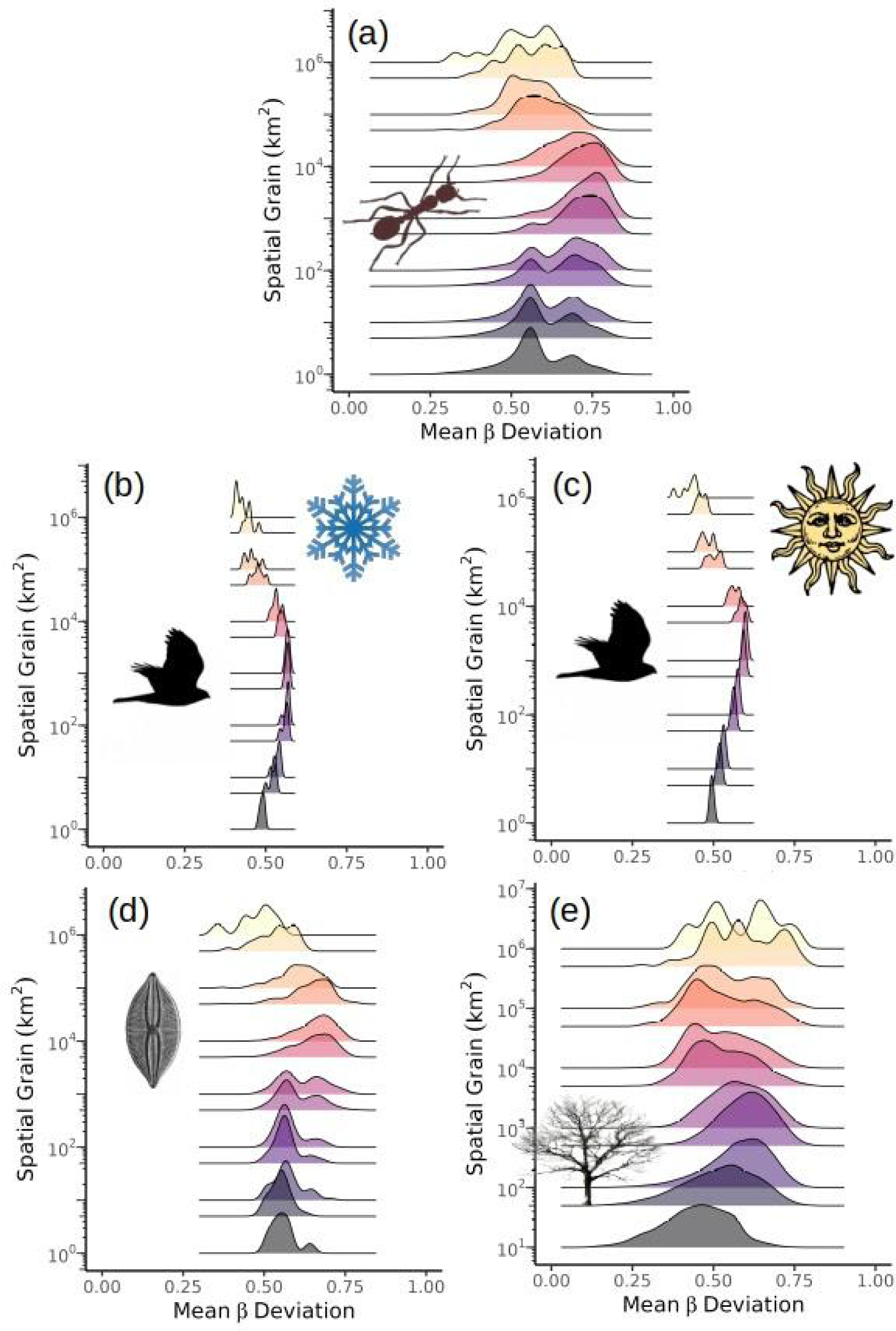
Data distributions for beta diversity, calculated as deviation from a null expectation (β_NBD_, Xing and He 2021) and potentially ranging 0-1. Normal data are best represented by means, and narrow variance generates narrow confidence intervals (see Fig. 2). (a) Ant β_NBD_ was most variable and multimodal at small and large spatial grains. Bird β_NBD_ in both (b) winter and (c) summer was relatively narrow overall but especially suited to means at peak β_NBD_. (c) Similar to ants, diatoms were best represented by means at peak values. (d) Trees data were multimodal at greatest spatial grains. Note log scale for spatial grain. Fig 1.

In all cases, spatial grains with peak β_NBD_ obtained data distributions most appropriately described by a mean and confidence intervals, whereas other spatial grains represented more problematic distributions. Thus, distributions of mean β_NBD_ (Fig. 3) validated inferences about peak mean β_NBD_ values at intermediate spatial grains across varied clades analyzed here.

To be clear, β deviation based on spatial homogeneity (β_DEV_) did increase with sample size as expected, but β_NBD_ did not. Instead, β_NBD_ consistently peaked at intermediate grain sizes that corresponded to intermediate numbers of samples, hexagons, species, and neighboring hexagonal grid cells. Based on results above (Figs. 2 & 3) and additional results summarized here (details are in Supplementary Information), mean pairwise β_NBD_ was not a simple progressive function of sample size, whether estimated as number of hexagonal grids or size of those hexagons (because bigger hexagons include more observation locations). Also, mean pairwise β_NBD_ did not increase with raw sample size (i.e., number of sample points), neighborhood size or species richness (Supplementary Information). This detail is secondary to our main question here but is important to an alternative hypothesis that β simply scales with sample size (e.g., Fig. 1b). Moreover, peak β_NBD_ occurred at different spatial scales among clades despite having identical base hexagonal maps at each grain size. In all, multiple lines of evidence demonstrated that sample size effects were not important contributors to patterns in spatial scaling of mean pairwise β_NBD_. Also, β_NBD_ repeatedly represented a modest adjustment to β_OBS_ whereas β_DEV_ represented substantial changes relative to β_OBS_.

## Discussion

Results here help address decades of confusion about the conceptual reality and practical value of ecological communities, as well as beta diversity (β) and spatial scaling (Whittaker 1972, McIntosh 1986, Wilson 1991, Lawton 1999, Ricklefs 2008, Roughgarden 2009, Vellend 2010, Tuomisto 2010, Anderson et al. 2011, Chave 2013). The community concept has been debated almost since its origin (McIntosh 1986, Wilson 1991, Lawton 1999, Ricklefs 2008, Roughgarden 2009). Ecology and biogeography have nonetheless progressed, often by ignoring this problem. At the same time, scale became widely recognized as fundamental to understand natural processes and patterns (Chave 2013), and β emerged (with its many versions; Tuomisto 2010, Anderson et al. 2011) as an important measure in ecology and biogeography. In that context, ecology has (in part) expanded toward biogeographical spatial scales (Lawton 1999, Leibold et al. 2004, Roughgarden 2009, Jenkins and Ricklefs 2011). Results here support this research direction by showing empirical differences between adjacent communities are most apparent at regional scales (hundreds to tens of thousands of km^2^), consistent with the regional community concept (Ricklefs 2008) for clades as different as ants, birds, diatoms and trees.

Work here was based on the premise that an entity is best measured by its own properties, and thus β (rather than external features; habitat, etc.) can indicate a spatial scale that most clearly differentiates between neighboring communities. We obtained empirical *gradients* of β across spatial grains, which were free to form inherent shapes and where any peak values represented spatial scales of the maximal differences between adjoining communities.

Below we elaborate on five extracted themes. First, long-standing work in remote sensing (e.g., Woodcock and Strahler 1987, Ming et al. 2011, Mutowo et al. 2023) to optimize spatial resolution of images provides lessons for how biogeography may improve understanding of spatiotemporal patterns. Secondly, results here indicate β_NBD_ has great value for analyses at biogeographical scales, where spatial heterogeneity is expected to exert strong effects on biodiversity patterns. Importantly, results here were broadly consistent across the tree of life and among rather different data origins, suggesting generality in the shape of β spatial scaling (and thus the regional community concept). Moreover, data for birds and trees indicate spatial scaling of β_NBD_ is inter-annually repeatable and (for birds) sensitive to seasonal conditions, suggesting common bases for β_NBD_ spatial scaling. Finally, results may help better understand diversity responses to land use and climate change, and help steer regional collaborations in conservation biology.

To evaluate spatial scaling of β, we adopted logic used to optimize spatial resolution in remote sensing data (Woodcock and Strahler 1987, Ming et al. 2011, Mutowo et al. 2023), which has wide relevance in spatial ecology and biogeography. Multiple other spatial patterns key to biogeography may also exhibit similar spatial scaling (i.e., peak values at some spatial grains), but that will be unknown if multiple spatial grains are not first compared before analyses are conducted. Our choice of biogeographical spatial grain should be based on maximal signal, rather than other criteria (e.g., space filling, or visual appeal). If so, ecological and biogeographical concepts tested so far at varied spatial grains and then debated may gain better resolution.

While β is a fundamental measure of ecological diversity patterns (Whittaker 1972), it has a long history of disputes spanning its metrics and concepts. That history largely coincided with debate on the veracity of ecological communities. Work to organize β (e.g., Tuomisto 2010, Anderson et al. 2011) made it more clearly understood, including the use of beta deviations from null expectations (β_DEV_, Kraft et al. 2011) that led to the recent update for measuring deviation of β from null expectations based on spatial heterogeneity (β_NBD_; Xing and He 2021) used here. We showed that β_NBD_ moderately adjusted observed β values (β_OBS_) and did not introduce sample size effects, whereas β_DEV_ greatly adjusted β_OBS_ and was subject to sample size effects. Thus, results here among very different clades and data sources support the value of β_NBD_ for future research on β diversity.

Very different clades (and data origins) obtained a remarkably consistent *general* shape for β spatial scaling here (yes, differences existed in the details). If this general pattern holds among other clades across the tree of life and different data sources, some small generality is enticingly possible for the spatial scaling of biodiversity patterns and the regional community concept. What processes might cause such general patterns? Much of our collective understanding of ecological processes is necessarily obtained at local scales much finer than those used here, related to historical, logistical limits on data collection. We stress that results here do not disavow local-scale research, which is vital to understand mechanisms of biotic and abiotic interactions and provided data amassed here and then integrated into regional spatial units. However, results here support efforts to better understand how myriad local processes (i.e., abiotic and biotic interactions affecting individuals or demes) may cascade up to cause regional patterns, and to compare those local processes to regional processes (Ricklefs and Schluter 1993, Ricklefs 2008, D’Amen et al. 2017). In this theme, it must also be remembered that results here were about β diversity; local and/or regional processes on individuals, demes, or a species must be general enough to translate to β diversity to be detected here.

Results here may also be informative for the dispersal scaling of communities, where potential long-range dispersal of microscopic organisms has long been debated as causing greater similarity than for macroscopic organisms (Fontaneto 2011). At first glance, spatial scaling of β_NBD_ for diatoms appeared to support long-range dispersal scaling of microbial diversity.

However, community size of lake diatoms roughly matched the average sampled lake density (1071 sampled lakes / 8.08 x 10^6^ km^2^ in the contiguous US = 1 lake / 1.3 × 10^4^ km^2^), and the majority (52%) of species were recorded in ≤3 lakes. Thus, diatom results here are consistent with a diatom community being bounded by one lake shore, on average. Spatial scaling of diatom β_NBD_ may actually represent the average dispersion of sampled discrete diatom habitats, in contrast to intergrading terrestrial habitats here for ants, birds, and trees. If so, other microbial communities (e.g., soils) may be evaluated with the approach here to further evaluate dispersal scaling of microbial community structure.

Results for birds and trees indicate spatial scaling of β_NBD_ is inter-annually repeatable and (for birds) sensitive to seasonal conditions. Given that birds and trees differ so greatly in life histories and mobility, this commonality suggests community assembly processes are a strong and common basis for β_NBD_ spatial scaling. Future work on the spatial scales of those processes may help explain the regional scales observed here.

Interestingly, ants and summer birds had roughly similar regional community scales, perhaps related to similar scales of factors that control distributions of birds and founding ant queens that establish colonies (Helms et al. 2016). Finally, differences in bird β_NBD_ scaling between summer and winter indicate that the approach here is sensitive to temporal conditions across substantial extent (contiguous US). Given this pattern, changes in β_NBD_ scaling over longer terms may represent effects of changing climate and/or land cover.

Finally, our results bear fundamental implications for basic and applied ecological disciplines. A regional community has been interpreted as the scope of an entire metacommunity (Jenkins and Ricklefs 2011). If that is the case, then results here represent the metacommunity in aggregate and identify its potential size. However, a subsequent definition of internal, local communities would remain a problem. Alternatively, if a single community is ∼10^3^ km^2^, then a metacommunity is vast. Metacommunity ecology needs to grapple with its definition of local communities relative to evidence here for regional communities. Additionally, results here indicate that regional units of study (i.e., tuned for the taxon of interest) are important to understand diversity responses to land use and climate change, as well as units for efforts to conserve biodiversity (Whittaker et al. 2005). We see results as adding importance to work toward corridors and networks among conserved sites (e.g., Rouget et al. 2006), and to help researchers consider spatial scales to use when studying diversity responses to land use and climate change. For example, a study to evaluate changes due to land use and climate change in rastered units of study may first need to optimize spatial grain to obtain maximal resolution before evaluating changes.

## Acknowledgements

We thank the Cornell Lab of Ornithology, the GABI network, the US Forest Service, and the USEPA for making many data publicly-available and hope results here help validate that work. We also thank the Ying Family Fund for continuing support.

## Biosketch

The authors started working and laughing together while at UCF, and have continued to do so despite the greater miles between them since. All are interested in macroscale ecological patterns and the processes that drive them.

## Data Availability

all data used here are already publicly-available and sources are listed.

## Funding

This research was conducted without specific funding.

## Conflict of Interest

All authors declare that there are no competing interests.

## Notes

### Competing Interest Statement

The authors have declared no competing interest.

