## Supplemental figures for "Spatial scaling of beta diversity supports the regional community concept for clades as different as ants, birds, diatoms, and trees"

**Supplementary Materials**

Xing & He (2021) demonstrated that $\beta_{DEV}$ scaled with sample size, according to a power law (see their Fig. 5), but did not test for similar effects on $\beta_{NBD}$. Similar to their results, $\beta_{DEV}$ repeatedly showed sample size effects in our analyses, whereas $\beta_{OBS}$ and $\beta_{BND}$ did not (Figs. S.1-S.3). We evaluated sample size effects in three ways: using simple number of samples (i.e., georeferenced observation points); species richness (S); and neighborhood size (M) of hexagonal grids, where >3 grids were used in calculations (data for 2 grids are depicted but gray to indicate neighborhood size = 2 was not used). We infer $\beta_{DEV}$ introduces sample effects not exhibited by $\beta_{OBS}$ and $\beta_{BND}$ (Tuomisto 2010, Xing and He 2021). Also, our use pairwise β calculations between each central hexagon and its neighbors effectively standardized neighborhood effects across spatial grains.

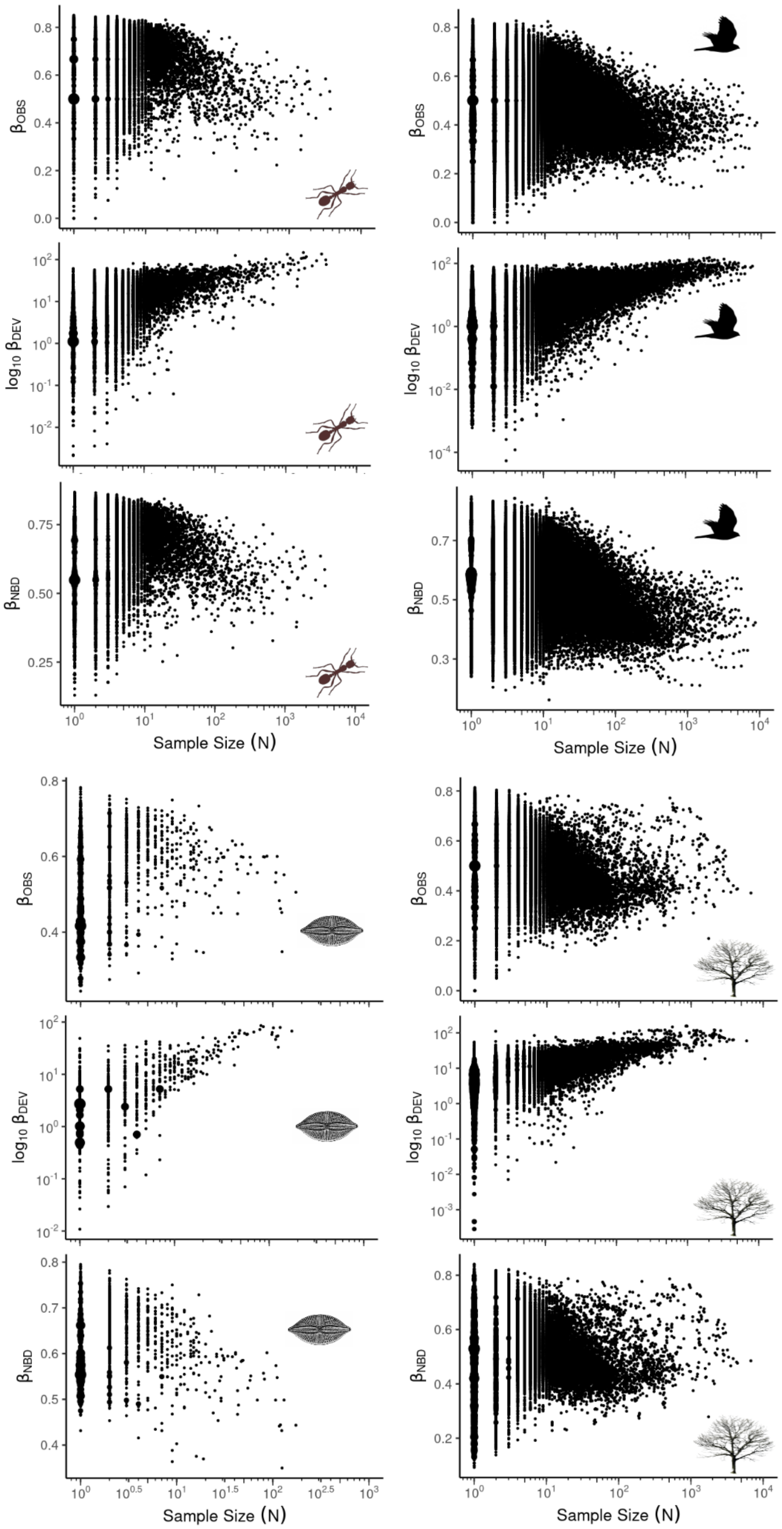
$\beta_{OBS}$
$\log_{10} \beta_{DEV}$
$\beta_{NBD}$
Sample Size (N)

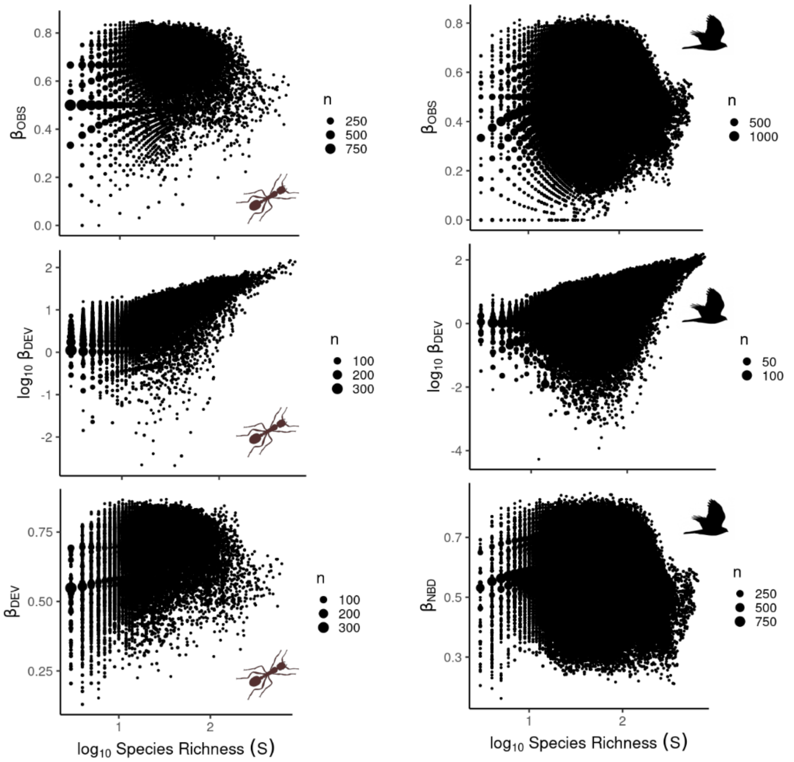
βOBS
n
250
500
750
log10 βDEV
n
100
200
300
βDEV
n
100
200
300
log10 Species Richness (S)
βOBS
n
500
1000
log10 βDEV
n
50
100
βNBD
n
250
500
750
log10 Species Richness (S)

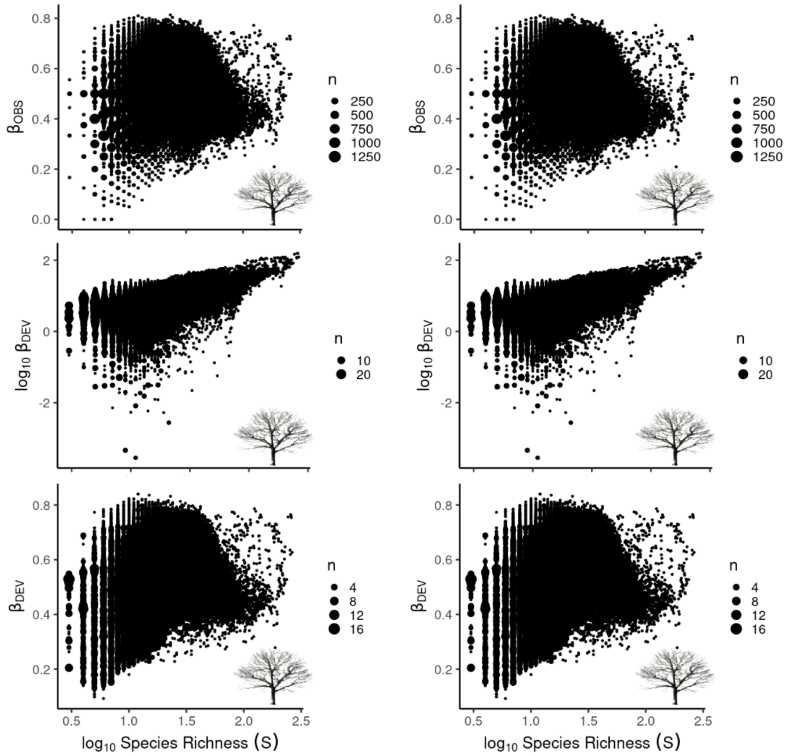
βOBS
n
250
500
750
1000
1250
log10 βDEV
n
10
20
βDEV
n
4
8
12
16
log10 Species Richness (S)
βOBS
n
250
500
750
1000
1250
log10 βDEV
n
10
20
βDEV
n
4
8
12
16
log10 Species Richness (S)

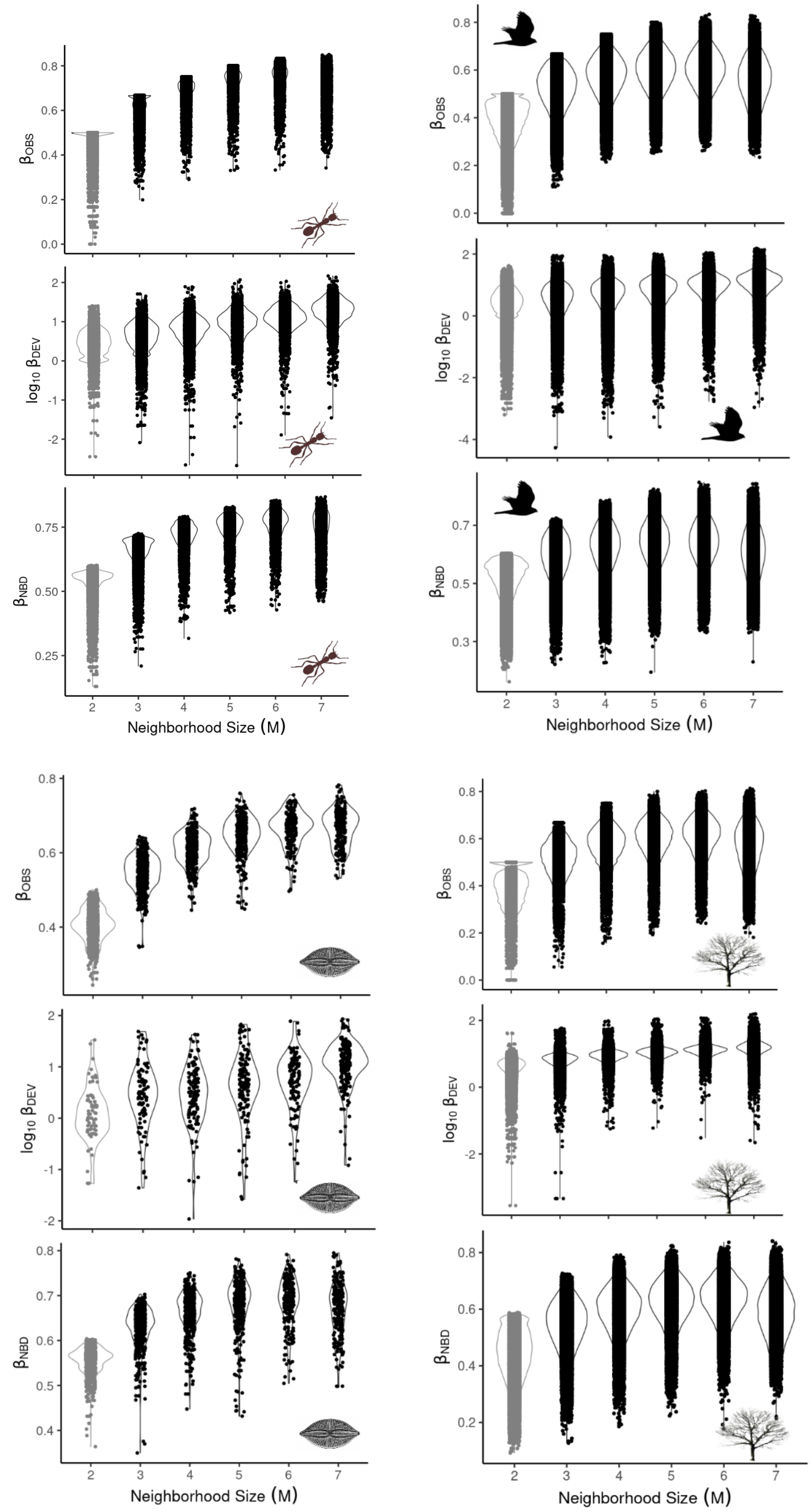

$\beta_{OBS}$
$\log_{10} \beta_{DEV}$
$\beta_{NBD}$
Neighborhood Size (M)
2
3
4
5
6
7
$\beta_{OBS}$
$\log_{10} \beta_{DEV}$
$\beta_{NBD}$
Neighborhood Size (M)
$\beta_{OBS}$
$\log_{10} \beta_{DEV}$
$\beta_{NBD}$
Neighborhood Size (M)
$\beta_{OBS}$
$\log_{10} \beta_{DEV}$
$\beta_{NBD}$
Neighborhood Size (M)
